# Active populations and growth of soil microorganisms are framed by mean annual precipitation in three California annual grasslands

**DOI:** 10.1101/2021.12.06.471491

**Authors:** Megan M. Foley, Steven J. Blazewicz, Karis J. McFarlane, Alex Greenlon, Michaela Hayer, Jeffrey A. Kimbrel, Benjamin J. Koch, Victoria Monsaint-Queeney, Keith Morrison, Ember Morrissey, Bruce A. Hungate, Jennifer Pett-Ridge

## Abstract

Earth system models project altered precipitation regimes across much of the globe; in California, the winter wet season is predicted to extend into spring, and the summer dry period to lengthen. How these precipitation trends will affect microbial traits and soil carbon (C) cycling is a key knowledge gap. Specifically, we do not have a mechanistic understanding of the linkages between soil moisture legacy effects, microbial population dynamics and soil C persistence. Using quantitative stable isotope probing (qSIP), we compared total and growing soil microbial communities across three California annual grasslands that span a rainfall gradient yet developed on similar parent material. We also assessed multiple edaphic variables, including the radiocarbon (^14^C) age of soil C, and found soil C turnover time increased with annual precipitation, but that soil microbes respired recently-fixed C regardless of site rainfall history. Samples were assayed in the wet season, when we expected environmental conditions would be most similar across sites. We hypothesized that growing communities would be more compositionally similar across the gradient than the total background microbiome. We also predicted that the long-term legacy effect of soil water limitation would be reflected in a lower community growth capacity at the driest site. We found that the proportion of the total community that was detected as growing was 28%, 48% and 58% at the wet, intermediate and dry sites, respectively. The composition of growing communities strongly resembled that of total communities, and growing communities were no more similar across the gradient than total communities, indicating a strong effect of climate of the structure of growing microbial communities. Members of three phyla, Acidobacteria, Actinobacteria, and Proteobacteria, were responsible for ∼79% of the cumulative ^18^O assimilation and 80% of all taxa that we defined as ‘growers’. Bacterial growth rates were low at the driest site relative to the intermediate and wettest sites. Reduced growth at the driest site was observed across major phyla, including the Actinobacteria, Acidobacteria, Bacteroidetes, Gemmatimonadetes and Proteobacteria. Microbial communities at the driest site displayed phylogenetic clustering, suggesting that climate history impacts microbial growth through environmental filtering for slow growing taxa. Taxonomic identity was a strong predictor of growth, such that the growth rates of a taxon at one site predicted its growth rates at the others. This cross-site coherence in growth is likely a consequence of genetically determined physiological traits, and is consistent with the idea that evolutionary history influences growth rate.

## 1. Introduction

Rainfall patterns and soil water content are crucial controllers of microbial population dynamics (growth and death) and Earth system models project major changes in the timing and intensity of precipitation events globally (Dore, 2005; Schimel, 2018). Since microorganisms mediate a wide range of ecosystem processes, including the accrual and persistence of soil carbon, understanding how the climate history of a site impacts microbial communities is essential. Precipitation is a key controller of ecosystems with Mediterranean climates (cool, wet winters and hot, dry summers) and it plays an equally important role in shaping microbial community structure and function (Angel et al., 2010; Maestre et al., 2015). Microorganisms in Mediterranean ecosystems must withstand both direct physiological stress during prolonged periods of low soil moisture yet be able to compete for resources when seasonal rains return and plant growth resumes (Barnard et al., 2020).

Fluctuations between wet and dry seasons in Mediterranean climates can drive dramatic changes in soil microbial community structure, but most diversity studies rely on DNA sequencing which cannot distinguish genetic material from growing, dormant, and recently dead organisms (Cruz-Martínez et al., 2009; Barnard et al., 2015; Carini et al., 2017). These fractions of the microbial community –growing, dormant, and dead – contribute to soil processes differently (Sokol et al., 2022). Populations that are actively turning over (growing and dying) have a prominent role in biogeochemical processes since their growth requires uptake and transformation of environmental substrates, while their death deposits cellular materials into the soil-mineral matrix where it may be subsequently recycled into living biomass or persist as soil organic matter (Liang et al., 2017). It widely accepted that microbial growth is a major factor in the formation of soil carbon (Bradford et al., 2013). The factors controlling which microbial taxa grow in soils and the timing of their growth are not well understood, but active communities have a heightened sensitivity to changes in environmental conditions compared to the total community (Angel et al., 2013; Baubin et al., 2019). In ecosystems with Mediterranean climates, strong seasonal changes in soil water may act as an important determinant of which taxa grow and when.

Soil water also strongly influences microbial function. Historic exposure to low soil water potential can have a lasting effect on microbial respiration, enzyme activity, and carbon use efficiency (Averill et al., 2016; Nijs et al., 2019; Hawkes et al., 2020), and is likely to influence growth as well. Repeated exposure to limited soil moisture may favor traits that confer tolerance to water stress and select against traits that enhance organisms’ ability to grow quickly or efficiently (Evans and Wallenstein, 2014; Malik et al., 2020a, 2020b). For example, the production of extracellular polymeric substances (EPS) can be a substantial carbon sink for bacteria growing under water stress, and osmolyte regulation in response to water stress can consume both nutrients and energy (Flemming and Wingender, 2001; Sandhya and Ali, 2015). In bacterial isolates, moisture stress has been shown to drive a tradeoff between EPS production and the length of lag phase, pointing to the relevance of growth rate in water stress acclimation (Lennon et al., 2012). While such traits provide a clear advantage during chronic water stress, there are likely biochemical costs involved in maintaining them. Thus, microbial taxa from soils routinely exposed to low soil water potentials may have acquired traits that lead to slow growth rates even in the absence of immediate water stress. At the community level, the mechanism underlying changes in growth rates may include a shift in microbial community composition, a change in the physiology of microbial taxa, or both (Evans and Wallenstein, 2014).

Climatic history can shape microbial function via shifts in composition or changes in microbial physiology. In this study, we compared total and growing microbial communities and soil characteristics across three California annual grasslands with Mediterranean-type climates that span a rainfall gradient and developed upon similar parent materials. First, we characterized the soils to understand how diverging precipitation regimes have shaped the physiochemical environment that soil microbes experience within each ecosystem. We also analyzed the soils’ radiocarbon (^14^C) age and ^14^C of CO_2_ to investigate how soil C persistence varies along the gradient. We then compared the structure of communities inferred from 16S rRNA sequencing (the “total community”) against the communities of growing microorganisms identified with quantitative stable isotope probing (qSIP) (the “growing community”) at each site during the wet season, when soil water was not limiting (Hungate et al., 2015; Koch et al., 2018). Hypothesizing that growing community structure is more strongly influenced by recent environmental conditions than climate history, we expected that similarity in soil water content during the weeks preceding sampling would result in growing communities that were compositionally more similar to each other than total communities. Lastly, we quantitatively analyzed patterns of microbial population growth using qSIP. We hypothesized that growth would be slowest in the driest site even during the wet season, reflecting a legacy effect of low soil moisture on microbial growth. We predicted that shifts in composition, likely due to habitat filtering for slow-growing drought tolerant taxa, would be evidenced by phylogenetic clustering in community composition (detected via net relatedness index and nearest taxon index).

## 2. Materials and Methods

### 2.1 Site description and sample collection

We characterized soil chemistry and bacterial and archaeal microbial communities in three coastal California annual grasslands with Mediterranean-type climates. The sites span a significant rainfall gradient (388 mm yr^-1^ to 2833 mm yr^-1^), yet have similar plant communities and developed on coastal terraces with similar parent material (Table 1). The wettest site, Angelo Coast Range Reserve, lies upstream of the headwaters of the South Fork Eel River where indigenous peoples including the Cahto, Pomo, Wailaki, Yuki, Weott, and Sinkyone historically held territory (Power et al., 2015). Annual grasslands at Angelo intersperse mixed-oak woodlands and old-growth conifer forests that occur on strath terraces which formed as the Eel River cut through the soft coastal sediment (Seidl et al., 1992; Sullivan et al., 2016). The intermediate site, Hopland Research and Extension Center, sits in the foothills of the Mayacamas Mountains and encompasses topographically rugged rangeland on territory historically occupied by the Pomo Nation. The driest site, Sedgwick Reserve, spans the Santa Ynez Valley and San Rafael Mountains and is located on land where the Chumash people have historically lived.

**Table 1:** Characteristics of three California annual grasslands examined in this study.

|  | **Angelo Coast Range**<br>**Reserve** | **Hopland Research and**<br>**Extension Center** | **Sedgwick Reserve** |
| --- | --- | --- | --- |
| Location | 39°44.25`N, 123°37.82`W | 39°00.11`N, 123°04.18`W | 34°42.70`N, 120°2.33`W |
| Elevation (m) | 475 | 180 | 360 |
| MAT max/min (°C) | 20/6 | 23/7 | 24/7 |
| MAP (mm yr<sup>-1</sup>) | 2160 | 956 | 383 |
| Parent Material | Argillite/Sandstone<br>interbedding | Sandstone, Shale, Graywacke,<br>Schist | Alluvium, Calcareous shale,<br>Acid sandstone |
| Soil Order | Alfisol | Alfisol | Mollisol |
| Soil Series | Holohan-Hollowtree-Casabonne complex | Witherell-Squawrock complex | Botella |
| Dominant<br>vegetation | *Aira spp.*, *Bromus spp.*,<br>*Briza spp.* | *Avena spp.*, *Bromus spp.*,<br>*Erodium spp.*, *Festuca spp.* | *Avena spp.*, *Bromus spp.* |

The three sites developed on similarly-aged sedimentary rock, primarily sandstone and shale from the Franciscan formation (Table 1). The parent material at Sedgwick also consists of alluvium from the Paso Robles formation which contains Franciscan mélange and Monterey shale (Homyak et al., 2018). Angelo soils are Ultic Haploxeralfs of the Hologan-Hollowtree-Casabonne complex (Berhe et al., 2012), Hopland soils are Typic Haploxeralfs of the Witherall-Squawrock complex (Fossum et al., 2021), and Sedgwick soils are Pachic Argixerolls belonging to the Botella series (Homyak et al., 2018). These taxonomies depict three naturally fertile soils (Angelo and Hopland are Alfisols while Sedgwick is a Mollisol) that experience a moderately dry, Mediterranean-type climate (Angelo and Hopland are Xeralfs while Sedgwick is Xeroll). Naturalized non-native annual grasses dominate vegetation at the three sites. *Avena spp.* and *Bromus spp.* dominate at Hopland and Sedgwick (Michael Barbour et al., 2007; Leitner et al., 2017). Vegetation at Angelo is dominated by non-native annual grasses including *Aira caryophyllea, Bromus hordeaceuous, Briza minor* and *Avena spp.* and by annual and perennial forbs interspersed with the native perennial grass *Danthonia californica* (Kotanen, 2004; Berhe et al., 2012; Sullivan et al., 2016) (Table 1).

We collected soils in February-March 2018. At each site, surface vegetation was removed and three replicate soil cores (10 cm x 5 cm) were collected 1 m apart for chemical characterization, radiocarbon analyses, and quantitative stable isotope probing. Soil cores were transported on ice to Lawrence Livermore National Laboratory where they were homogenized by hand, stones and large roots were removed, and portions were dried at room temperature, stored at 4 °C, or frozen at -80 °C for different downstream analyses. All analyses reported here were conducted on surface soils (0-10 cm). We measured gravimetric soil water content by drying soils to a constant weight at 105°C. We retrieved daily mean air temperature and total precipitation data from nearby climate stations and on-site records for the calendar year and thirty-day period preceding sample collection at each site.

### 2.2 Soil physical and chemical characterization

Soil chemical and physical characterization was conducted at the Oregon State University Soil Health Laboratory. Bulk C and nitrogen (N) concentrations were quantified using an Elementar vario MACRO cube elemental analyzer through dry combustion, separation of gaseous species, and thermal conductivity detection. Trace elements were quantified on an Agilent 5110 ICP-OES after soils were extracted with 0.002M barium chloride. Soil texture was determined by the hydrometer method. To measure pH, 4 g of soil was suspended in 8 mL of 0.01 M CaCl_2_ and equilibrated for 1 hour prior to analysis (Minasny et al., 2011). Soil pH was measured using a Mettler Toledo Seven Compact pH/Ion meter. Additional soil cores (0-5 cm) were collected in June of 2019 to construct soil water retention curves for the three sites. Soil cores were also collected from the top 8-15 cm of soil to calculate bulk density for the retention curves. Intact soil cores were prepared and analyzed using Hyprop and WP4C instruments in accordance with manufacturer’s instructions (Pertassek et al., 2011; Meter Group, 2019).

### 2.3 Quantitative x-ray diffraction

Quantitative x-ray diffraction (qXRD) was performed at Lawrence Livermore National Laboratory as previously described (Fossum et al., 2021). Samples for qXRD were crushed and passed through a 500 µm sieve. 3 g of soil was then ground for 5 minutes with 15 mL of methanol using a MCrone mill. The ground samples were transferred to a plastic tray, air dried, and homogenized for 3 min using a vortex mixer with 10 mm plastic beads (Bakker et al., 2018). The soil powders were side loaded into XRD sample mounts. All powdered samples were analyzed on a Bruker D8 advance XRD. The samples were scanned from 3° to 65° 2θ at 0.011° steps with a 5 second per step count time. Quantitative mineralogy was determined using the BGMN Rietveld refinement and the Profex interface software (Doebelin and Kleeberg, 2015). XRD patterns were refined to fit crystal unit cell parameters, size, site occupancy, and preferred orientation.

### 2.4 Natural abundance stable isotopes and radiocarbon analysis of bulk soils and incubations

For each core, subsamples of dried homogenized soil were analyzed for δ^13^C and δ ^15^N at the Center for Stable Isotope Biogeochemistry at the University of California, Berkeley (IsoPrime100 mass spectrometer) or prepared for bulk soil radiocarbon measurement by sealed-tube combustion to CO_2_ in the presence of CuO and Ag. A subsample of fresh soil from each core was incubated in the lab to measure Δ^14^CO_2_ of microbially respired carbon. After removing roots from soils by hand picking, soils were left undisturbed for one week at room temperature to allow any remaining roots to senesce. For each core, approximately 150 g dry mass-equivalent soil was weighed into 150 mL beakers and placed inside quart-sized mason jars with stopcock-fitted lids. Soils were pre-incubated for one day before incubation jars were sealed and headspaces flushed with CO_2_-free air. Soils were then incubated at room temperature until enough CO_2_ had accumulated for ^14^C measurement (9-11 days). At the end of the incubation period, headspace CO_2_ was cryogenically purified for radiocarbon analysis. Aliquots of purified CO_2_ were analyzed for δ^13^C at the UC Davis Stable Isotope Laboratory (GVI Optima Stable Isotope Ratio Mass Spectrometer).

CO_2_ from both combusted bulk soil and incubation jars was purified cryogenically at the Center for Accelerator Mass Spectrometry (CAMS) at Lawrence Livermore National Laboratory using a vacuum line before being reduced to graphite on iron powder in the presence of H_2_ (Vogel et al., 1984). Radiocarbon abundance in graphitized samples was determined on the FN Van de Graaff Accelerator Mass Spectrometer at Lawrence Livermore National Laboratory’s Center for Accelerator Mass Spectrometer. Radiocarbon values are reported in Δ^14^C notation corrected for mass-dependent fractionation using measured δ^13^C values (Stuiver and Polach, 1977, p. 14).

### 2.5 ^18^O-water quantitative stable isotope probing (qSIP)

For qSIP incubations, 5 g soil from each field replicate was transferred to 15 ml Nalgene flatbottom vials. Soils were lightly air dried in a laminar flow hood at room temperature for 24 hours prior to isotope addition. This drying is necessary to allow for sufficient enrichment of the soil water with the isotope label while also maintaining soil moisture reasonably similar to field moisture during the qSIP incubations. One milliliter of isotopically enriched water (98.15 at% ^18^O-H_2_O) or natural abundance water (as a control) was pipetted onto the soil slowly and evenly and then gently mixed with the pipette tip (raising soil moisture content by 3-7%). After the water addition, vials were immediately sealed inside 500 ml mason jars and incubated at room temperature in the dark for 8 days. At the end of the incubation, soils were frozen in liquid nitrogen and then stored at -80°C.

DNA was extracted from all soil samples using a modified protocol adapted from Barnard et al. (2015). Three replicate extractions were conducted for each sample and then replicate DNA extracts were combined. For each extraction, soil (0.4 g +/-0.001 g soil) was added to 2 ml Lysing Matrix E tube (MP Biomedicals) and extracted twice as follows. 500 µl extraction buffer (5% CTAB, 0.5 M NaCl, 240 mM K_2_HPO_4_, pH 8.0) and 500 µl 25:24:1 phenol:chloroform:isoamyl alcohol were added before shaking (FastPrep24, MP Biomedicals: 30 s, 5.5 m ^s-1^). After centrifugation (16,100 x g, 5 min), residual phenol was removed using pre-spun 2 ml Phase Lock Gel tubes (5 Prime, Gaithersburg, MD, USA) with an equal volume of 24:1 chloroform:isoamyl alcohol, mixed and centrifuged (16,100 x g, 2 min). The aqueous phases from both extractions were pooled, mixed with 7 µl RNAase (10 mg/ml), mixed by inverting, and incubated at 50 °C for 10 min. 335 µL 7.5 M NH4+ acetate was added, mixed by inverting, incubated (4 °C, 1 h). and centrifuged (16,100 x g, 15 min. The supernatant was transferred to a new 1.7 ml tube and 1 µl Glycoblue (15 mg/ml) and 1 ml 40% PEG 6000 in 1.6 M NaCl was added, mixed by vortex, and incubated at room temperature in the dark (2 h). After centrifugation (16,100 x g, 20 min), the pellet was rinsed with 1 ml ice-cold 70% ethanol, air-dried, resuspended in 30 µl 1xTE and stored at -80 °C.

Samples were subjected to a cesium chloride density gradient formed by physical density separation via ultracentrifuge as previously described with minor modifications (Blazewicz et al., 2014; Sieradzki et al., 2020). For each sample, 5 µg of DNA in 150 µL 1xTE was mixed with 1.00 mL gradient buffer, and 4.60 mL CsCl stock (1.885 g mL^-1^) with a final average density of 1.730 g mL^-1^. Samples were loaded into 5.2 mL ultracentrifuge tubes and spun at 20 °C for 108 hours at 176,284 RCF_avg_ in a Beckman Coulter Optima XE-90 ultracentrifuge using a VTi65.2 rotor. Automated sample fractionation was performed using Lawrence Livermore National Laboratory’s high-throughput SIP (‘HT-SIP’) pipeline, which automates fractionation and clean-up tasks for the density gradient SIP protocol. Ultracentrifuge tube contents were fractionated into 36 fractions (∼200 µL each) using an Agilent Technologies 1260 isocratic pump delivering water at 0.25 mL min^-1^ through a 25G needle inserted through the top of the ultracentrifuge tube. Each tube was mounted in a Beckman Coulter fraction recovery system with a side port needle inserted through the bottom. The side port needle was routed to an Agilent 1260 Infinity fraction collector. Fractions were collected in 96-well deep well plates. The density of each fraction was then measured using a Reichart AR200 digital refractometer fitted with a prism covering to facilitate measurement from 5 µL, as previously described (Buckley et al., 2007). DNA in each fraction was purified and concentrated using a Hamilton Microlab Star liquid handling system programmed to automate previously described glycogen/PEG precipitations (Neufeld et al., 2007). Washed DNA pellets were suspended in 40 µL of 1xTE and the DNA concentration of each fraction was quantified using a PicoGreen fluorescence assay. The fractions for each sample were binned into 9 groups based on density (1.6900-1.7099 g/ml, 1.7100-1.7149 g/ml, 1.7150-1.7199 g/ml, 1.7200-1.7249 g/ml, 1.7250-1.7299 g/ml, 1.7300-1.7349 g/ml, 1.7350-1.7399 g/ml, 1.7400-1.7468 g/ml, 1.7469-1.7720 g/ml), and fractions within a binned group were combined and sequenced.

For 16S rRNA gene amplicon sequencing, non-fractionated DNA as well as density fractionated DNA was amplified in triplicate 10-uL reactions using primers 515 F and 806 R (Apprill et al., 2015; Parada et al., 2016). Each reaction contained 1uL sample and 9uL of Phusion Hot Start II High Fidelity master mix (Thermo Fisher Scientific) including 1.5mM additional MgCl_2_. PCR conditions were 95 C for 2 min followed by 20 cycles of 95 C for 30 S, 64.5 C for 30 S, and 72 C for 1 min. The triplicate PCR products were then pooled and diluted 10X and used as a template in a subsequent dual indexing reaction that used the same primers including the Illumina flowcell adaptor sequences and 8-nucleotide Golay barcodes (15 cycles identical to initial amplification conditions). Resulting amplicons were purified with AMPure XP magnetic beads (Beckman Coulter) and quantified with a PicoGreen assay on a BioTek Synergy HT plate reader. Samples were pooled at equivalent concentrations, purified with the AMPure XP beads, and quantified using the KAPA Sybr Fast qPCR kit (Kapa Biosciences). A total of 9 unfractionated, and 162 fraction libraries were sequenced on an Illumina MiSeq instrument at Northern Arizona University’s Genetics Core Facility using a 300-cycle v2 reagent kit. Sequence data and sample metadata have been deposited in the NCBI Sequence Read Archive under the accession number PRJNA718849.

Paired-end 151 nt reads were filtered to remove phiX and other contaminants with bbduk v38.56 (default settings except k=31 and hdist=1) (Bushnell, 2014). Fastq files were then filtered/trimmed for quality (maxEE=5, truncQ=2) and used to generate amplicon sequence variants (ASVs) with DADA2 v1.10 and phyloseq v1.26 (McMurdie and Holmes, 2013; Callahan et al., 2016). Chimeric sequences were determined and removed using removeBimeraDenovo from DADA2. ASV taxonomy was determined using the RDP 16S rRNA gene database (training set 16) using RDP classifier v2.11, keeping classifications with greater than 50% confidence (Wang et al., 2007). A phylogenetic tree was built using Muscle v3.8.31 and FastTree v2.1.10 (Edgar, 2004; Price et al., 2010).

### 2.6 Quantitative stable isotope probing analysis

Quantitative stable isotope probing (qSIP) measures the change in buoyant density of a taxon’s DNA, or segment of DNA, due to assimilation of a substrate-derived heavy isotope. qSIP with ^18^O-H_2_O identifies actively growing microbial taxa since ^18^O is incorporated into the phosphodiester backbone of DNA during replication through the rapid exchange of oxygen between water and inorganic phosphate species (Schwartz, 2007). The amount of ^18^O incorporation into microbial DNA during an incubation with ^18^O-H_2_O thus approximates a microorganisms growth rate.

We quantified excess atom fraction (EAF) ^18^O of bacterial DNA following a modified version of the procedure described by Hungate et al. (2015) and Koch et al. (2018) using average DNA concentration to normalize the relative abundance of taxa within each density fraction (Papp et al., 2018a; Sieradzki et al., 2020) . We calculated median values and 95% confidence intervals for EAF ^18^O by bootstrapping (n=1000) across experimental replicates. qSIP analysis was limited to taxa that occurred in at least 2 (of 3) experimental replicates and in at least 2 (of 9) density fractions. These criteria were chosen to reduce the likelihood of falsely interpreting spurious density shifts as growth. Technical error associated with tube-level differences in CsCl density gradients was corrected as previously described (Morrissey et al., 2017). All qSIP calculations are available at https://github.com/mmf289.

When comparing values of ^18^O enrichment between sites, we chose to use the metric “fraction of maximum potential enrichment” or “(FME) ^18^O” to account for slight differences in the enrichment of the soil water during incubation. FME ^18^O was computed for each taxon as FME ^18^O = ^18^O-EAF_taxon_/average soil water ^18^O enrichment during eight-day incubation. For all analyses, ‘growing’ microbial taxa were identified as ASVs having 95% confidence intervals for EAF ^18^O exceeding zero. We refer to the community of growing microbial taxa at each site as the “growing microbial community”. We refer to the microbial community inferred from 16S rRNA marker gene sequencing of non-fractionated DNA at each site as the “total microbial community”.

### 2.7 Statistical analyses

All statistical analyses were conducted in R (R Core Team, 2021). Soil properties, relative abundances of minerals, and bulk and respired soil radiocarbon values were compared between sites using one-way ANOVA and Fisher’s LSD test. A Bonferroni correction was applied when multiple comparisons were made.

The relative abundances of taxa in total communities were measured from sequencing of unfractionated DNA samples. The relative abundances of taxa in growing communities were computed by removing counts from taxa that were not identified as growing via qSIP from unfractionated DNA samples and then re-computing relative abundance values. We computed the proportion of the total community that was growing by dividing the number of ASVs identified as growing by the total number of ASVs inferred from 16S rRNA sequencing at each site. We then used Bray-Curtis dissimilarity to assess between-site dissimilarity for both growing and total communities with vegan: Community Ecology Package (Wagner et al. 2019). Principal coordinate analysis (PCoA) was used to visualize community structure. We tested for differences in community composition between sites using ANOSIM on ranked Bray-Curtis dissimilarities. To test if the degree of community dissimilarity between sites differed for total versus growing microbial communities, we performed a t-test on mean Bray-Curtis dissimilarity for each pairwise comparison of sites between total and growing communities. A Bonferroni correction was applied to account for multiple comparisons.

We used one-way ANOVA and Fisher’s LSD to test for differences between sites in community, phylum, and family mean FME ^18^O and only included taxa that were identified as growing at each site in this analysis. A Bonferroni correction was applied when multiple comparisons were made. We also compared FME ^18^O of taxa that were uniquely growing at Sedgwick to the FME ^18^O of taxa that were growing at Sedgwick and at least one other site using a t-test. We conducted NRI and NTI analyses using the Picante package to determine phylogenetic relatedness of total microbial communities at each site.

We used linear regression to assess the relationship of taxon-specific FME ^18^O between sites and included only individual ASVs that were co-occurring in at least two sites. Phylogenetic signal analyses were used to test whether the growth (EAF ^18^O) of related organisms resembled each other more than would be expected by chance alone^1^. Phylogeny was built using the SILVA v128 tree using SATé-enabled phylogenetic placement (Janssen et al., 2018; Bolyen et al., 2019). We measured Blomberg’s K and Pagel’s λ using phytools (Revell, 2012) for taxon-specific EAF ^18^O values within each site and included all taxa in this analysis.

## 3. Results

### 3.1 Precipitation and air temperature trends prior to sample collection

We gathered precipitation and air temperature data to assess the environmental conditions at each site during the year and month prior to sample collection. The three sites form a gradient in MAP and MAT with precipitation decreasing and temperature increasing between sites in the order: Angelo, Hopland, and Sedgwick (Table 1). In the calendar year prior to sample collection, the wet season at Angelo and Hopland lasted until April. Precipitation remained scarce or absent during the summer months and returned again by November. In contrast, the wet season at Sedgwick occurred in only January and February and didn’t return until the following January (Supplementary Figure 1). In sum, Angelo was the wettest and coldest site and Sedgwick was hottest and driest site during the year preceding sample collection (Supplementary Table 1). During the 30 days leading up to sample collection, total precipitation differed between our sites by an average of only 46 mm and soil moisture of the field-collected soil varied modestly from 17% to 25% (Supplementary Tables 1 & 2). The average air temperature during the 30 days leading up to sample collection ranged from 3.76 °C at Angelo to 10.35 °C at Sedgwick.

### 3.2 Soil characterization

To understand how the range in precipitation regimes correlates with the physiochemical environment that soil microbes experience within each ecosystem, we characterized soil mineralogy via quantitative x-ray diffraction (qXRD), constructed soil water retention curves, and determined soil texture, total C and N, trace elements, pH, and effective cation exchange capacity. Soils at the three sites contain the same minerals in varying proportions (Supplementary Table 3). Of the non-clay minerals at the sites, quartz is proportionally most abundant by weight followed by plagioclase and K-feldspar. At the wet and intermediate sites, Angelo and Hopland, muscovite is the most abundant clay mineral followed by chlorite and kaolinite; at the dry site, Sedgwick, kaolinite is most abundant.

Sedgwick, the driest site, contained the highest proportion of clay particles (Table 2). Soil water retention curves show that Sedgwick soils also experience lower soil water potentials at higher moisture contents than the intermediate and wet sites (Supplementary Figure 1). Soil carbon and nitrogen were low at all sites and exhibited little variation. δ^15^N of bulk soil did not vary between sites and δ^13^C was slightly lower at Sedgwick than Hopland or Angelo. Soil C:N exhibited little variation between sites. pH decreased significantly with increasing MAP. Effective cation exchange capacity and most trace elements were highest in Sedgwick soil, with the exception of aluminum which was highest in the Angelo soil (Table 2).

**Table 2:** Physical and chemical characteristics of surface soils from three northern California 986 annual grasslands: Angelo, Hopland, and Sedgwick (mean ± standard error, n = 3). Differences 987 between site means were assessed by one-way ANOVAs and Fisher’s LSD. Bold text indicates 988 <u>ANOVA P-values less than 0.05, and BDL signifies values below detection limit.</u>

| | **Angelo**<br>Avg ± SE | **Hopland**<br>Avg ± SE | **Sedgwick**<br>Avg ± SE | **ANOVA**<br>*P* |
| --- | --- | --- | --- | --- |
| $\delta^{13}\text{C}$ Bulk Soil (‰) | -27.54 ± 0.1 | -27.64 ± 0.1 | -26.99 ± 0.2 | **0.04** |
| $\delta^{15}\text{N}$ Bulk Soil (‰) | 3.75 ± 0.4 | 4.23 ± 0.3 | 4.85 ± 0.2 | 0.10 |
| pH | 5.02 ± 0.02 | 5.55 ± 0.03 | 6.99 ± 0.03 | **<0.01** |
| Total base cations<br>(meq kg<sup>-1</sup> soil) | 10.49 ± 0.7 | 10.79 ± 0.7 | 27.61 ± 0.9 | **<0.01** |
| ECEC (meq kg<sup>-1</sup> soil) | 10.99 ± 0.4 | 10.84 ± 0.4 | 27.61 ± 0.5 | **<0.01** |
| % Sand | 28 ± 2 | 45 ± 1 | 38 ± 1 | **<0.01** |
| % Silt | 42 ± 1 | 36 ± 1 | 28 ± 2 | **<0.01** |
| % Clay | 27 ± 1 | 19 ± 1 | 34 ± 0 | **<0.01** |
| % C | 1.66 ± 0.3 | 1.39 ± 0.2 | 1.99 ± 0.1 | 0.22 |
| % N | 0.13 ± 0.02 | 0.11 ± 0.01 | 0.18 ± 0.01 | **0.03** |
| C:N | 12.19 ± 0.6 | 12.37 ± 0.2 | 11.04 ± 0.1 | 0.11 |
| Al (mg kg<sup>-1</sup>) | 44.77 ± 1.1 | 5.17 ± 0.4 | *BDL* | **<0.01** |
| Ca (mg kg<sup>-1</sup>) | 1455.44 ± 53.0 | 1350.63 ± 47.6 | 2380.67 ± 50.3 | **<0.01** |
| K (mg kg<sup>-1</sup>) | 98.98 ± 6.3 | 114.92 ± 13.9 | 390.39 ± 29.4 | **<0.01** |
| Mg (mg kg<sup>-1</sup>) | 353.39 ± 18.0 | 448.75 ± 14.9 | 1760.32 ± 38.3 | **<0.01** |
| Na (mg kg<sup>-1</sup>) | 3.44 ± 1.2 | *BDL* | 7.21 ± 1.2 | **0.03** |

### 3.3 Bulk soil and respired Δ^14^C

The Δ^14^C of bulk surface soils increased along the gradient from the wettest to the dry site, indicating that the average age of bulk soil C is oldest at Angelo, the site with the highest annual rainfall (Figure 1). Bulk soil Δ^14^C values were above modern at Sedgwick, indicating the presence of carbon associated with atmospheric weapons testing in the 20^th^ century, while the more depleted Δ^14^C values observed for soils from Angelo suggest the presence of older and more slowly cycling soil C at this site compared to the other sites. Δ^14^C at our intermediate rainfall site, Hopland, falls between these two extremes. For reference, turnover times— determined using a single-pool, steady-state, homogenous model (Torn et al., 2009)—were approximately 200, 400, and 700 years for Sedgwick, Hopland, and Angelo, respectively. In contrast, the Δ^14^C of respired CO_2_ was not significantly different for the three soils, indicating that microorganisms uniformly respired recently fixed C (with an average age of 7 years since fixation from the atmosphere), despite the differences in the average age of bulk soil carbon.

**Figure 1:**
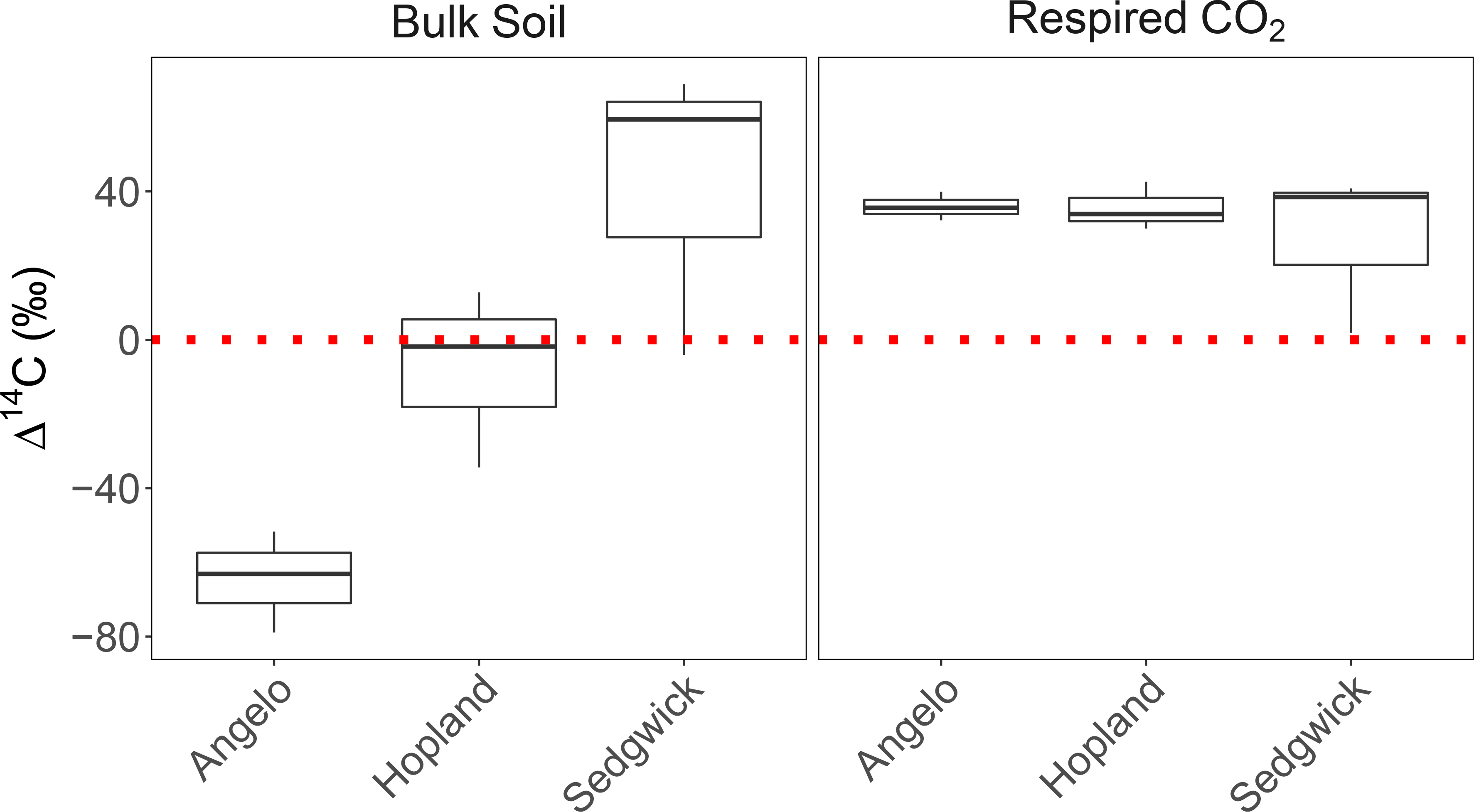
Δ^14^C (‰) of bulk soil and respired CO_2_ from three CA annual grasslands. The horizontal line represents atmospheric Δ^14^C in 2018 and serves as a reference for post-bomb enrichment. Differences between grassland site means (n=3) were assessed by one-way ANOVAs and Fisher’s LSD. Δ^14^C (‰) of bulk soils differed between sites (p<0.05, Fisher’s LSD) except Hopland did not differ from Sedgwick (p>0.05, Fisher’s LSD). Δ^14^C (‰) of respired CO_2_ did not differ between sites (p>0.05, Fisher’s LSD).

### 3.4 Structure of growing populations versus the total community

To test the hypothesis that growing communities would be compositionally more similar to each other than total communities, we analyzed the structure of growing and total communities using PCoA and pairwise comparisons of Bray-Curtis dissimilarity. The proportion of the total community that was detected as growing (ASVs having 95% confidence intervals for EAF ^18^O exceeding zero were considered growing) was 28%, 48% and 58% at the Angelo, Hopland and Sedgwick sites, respectively (Supplementary Table 4). Community structure differed by site for both the total and growing bacterial communities (Figure 2). Growing communities were not compositionally more similar to each other than total communities (Supplementary Figure 3). Sedgwick was the most dissimilar from the other sites, as quantified by mean pairwise Bray-Curtis Dissimilarity (Supplementary Figure 3).

**Figure 2:**
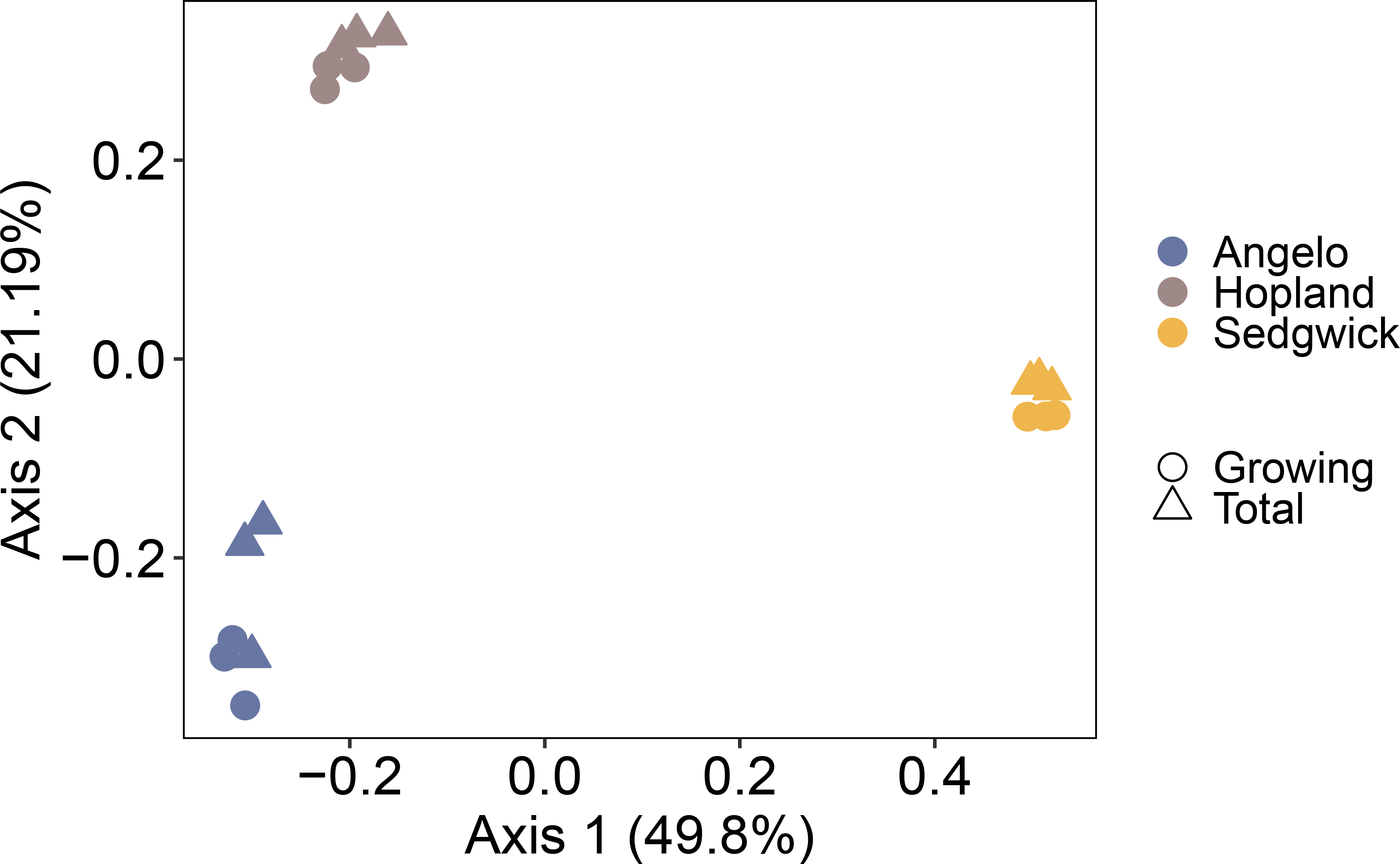
PCoA of Bray-Curtis dissimilarities for total (n=3) and growing (n=3) microbial communities assessed via 16S rRNA gene abundance and H_2_^18^O stable isotope probing in three CA annual grasslands, sampled during the winter wet season. Community composition varied significantly by site for the total and growing microbial communities (p < 0.01, ANOSIM).

More than 70% of 16S rRNA gene sequences in the total bacterial community at each site could be attributed to three phyla: Acidobacteria, Actinobacteria (dominant orders Rubrobacterales, and Solirubrobacterales), and Proteobacteria (dominant orders Burkholderiales, Sphingomonadales, and Rhizobiales). These phyla dominated growing communities as well, comprising more than 80% of all bacterial 16S rRNA gene sequences that were growing at each site and accounting for at least 79% of cumulative ^18^O assimilation in each site (Supplementary Figure 4). While these phyla accounted for a large proportion of total growth, there was no detectable relationship between the relative abundance of an individual taxon and its growth, meaning some low abundance taxa exhibited fast growth (Supplementary Figure 5).

### 3.5 Microbial growth along the precipitation gradient

Using qSIP, we analyzed patterns of microbial population growth to test the hypothesis that growth would be slowest in the historically driest site –even during the wet season— reflecting the climate history of the site. We used fraction of maximum potential enrichment (FME) ^18^O when comparing values of ^18^O enrichment between sites to account for small differences in the enrichment of the soil water during incubation. Site specific variation in microbial growth was present at the community, phylum, and family levels. The mean community FME ^18^O was significantly higher in Hopland and Angelo soils than in Sedgwick soils (Figure 3). These patterns, in general, also persisted across phyla and family levels: Actinobacteria and Bacteroidetes had a higher FME ^18^O in Hopland relative to Sedgwick soils, and Acidobacteria, Gemmatimonadetes, and Proteobacteria had higher FME ^18^O in both Angelo and Hopland soils (Figure 3). While the mean FME ^18^O of Comamonadaceae was highest in Sedgwick, this result was not statistically significant (P=0.33, ANOVA). No microbial families exhibited significantly faster growth in Sedgwick soils relative to the other sites (Supplementary Figure 6).

**Figure 3:**
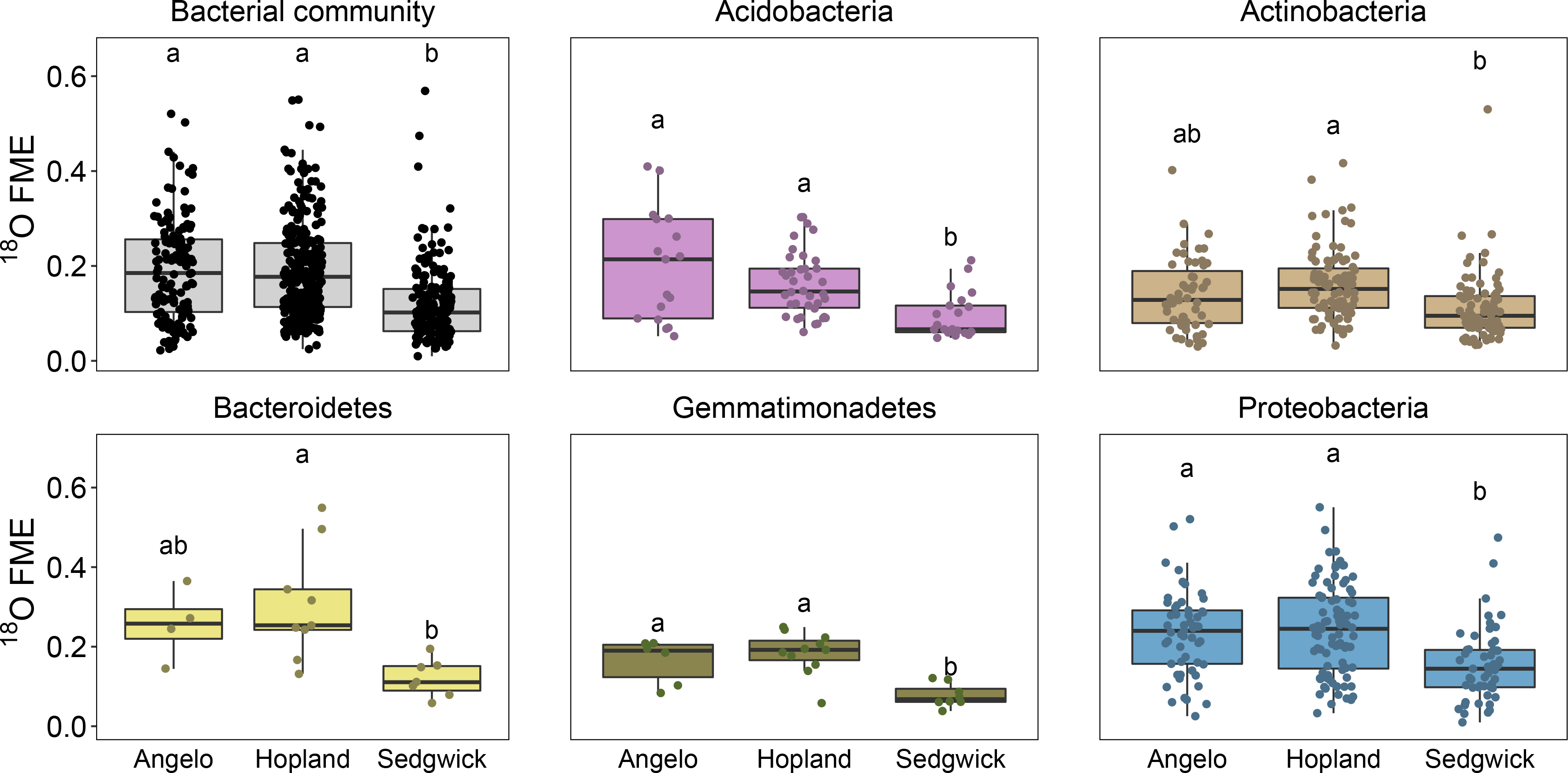
Community and phylum mean fraction of maximum enrichment ^18^O (FME) of soil microbial taxa that are growing (95% C.I. for excess atom fraction ^18^O > 0) at three CA annual grassland sites. Differences between grassland site means were assessed by one-way ANOVAs and Fisher’s LSD. Letters indicate significant differences between sites (p<0.05, Fisher’s LSD). FME ^18^O is computed as the excess atom fraction ^18^O of a taxon divided by the enrichment of the soil water during the qSIP incubation.

We calculated net relatedness index (NRI) and nearest taxon index (NTI) to assess whether reduced microbial growth at the driest site was driven by shifts in community composition (i.e. through habitat filtering for slow growing taxa). NRI and NTI analyses indicate phylogenetic clustering of total microbial communities at all three sites (Supplementary Table 6). We then grouped the taxa growing at Sedgwick according to whether they grew at Sedgwick alone or in at least one other site. We found that taxa that were unique to Sedgwick were slower growing than taxa that were growing in at least one other site (Figure 4).

**Figure 4:**
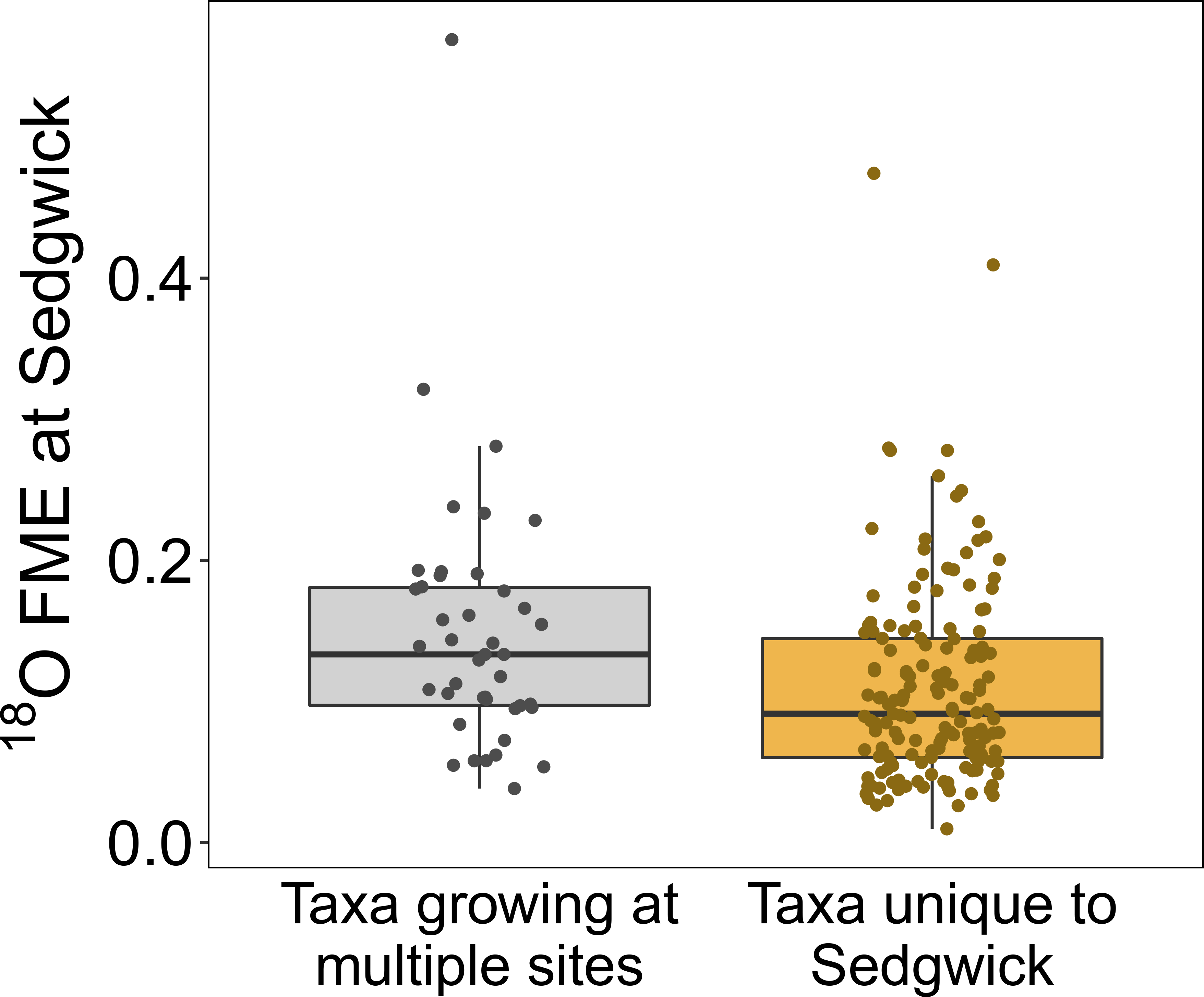
Fraction of maximum enrichment ^18^O (FME) of taxa growing in Sedgwick soil versus other annual grassland soils. Taxa plotted on the left were those growing in Sedgwick soils as well at least one other site. Taxa plotted on the right were found growing in Sedgwick soils alone. The FME ^18^O of taxa unique to Sedgwick soils is lower than that of taxa that grow in at least one other site (p-value = 0.01, t-test). FME ^18^O is computed as the excess atom fraction ^18^O of a taxon divided by the enrichment of the soil water during the qSIP incubation.

### 3.6 Evolutionary constraints on microbial growth

To better understand the factors driving variation in growth observed in our study, we used phylogenetic signal analyses to test whether taxon-specific growth was influenced by a microorganism’s evolutionary history. In all three grassland sites, we detected significant phylogenetic signals for taxon-specific growth, as determined by Pagel’s λ and Blomberg’s K (Supplementary Table 7). Stronger phylogenetic signals were observed in Angelo soil (λ = 0.82, p<0.001, K=0.30, p=0.001) and Hopland (λ = 0.85, p<0.001, K=0.10, p=0.004) than Sedgwick (λ = 0.68, p<0.001, K = 0.10, p=0.010). A significant phylogenetic signal indicates that the growth rates of closely related taxa resemble each other more than taxa drawn randomly from the same phylogenetic tree, however the values we observed are somewhat lower than expected under Brownian motion evolution (λ = 1 and K=1).

Given the evidence for a phylogenetic signal of in-situ microbial growth, we were curious to know if taxa would consistently perform with high or low activity regardless of the site, so we used linear regression to analyze taxon-specific growth rates across sites. For taxa that occurred at more than one site and were also significantly growing, their growth rate at one site was predictive of growth rates in another site (Figure 5). We compared ^18^O enrichment of bacterial ASVs that were present and growing in at least two sites and found positive, linear correlations of taxon-specific enrichment between sites. The growth rate of a bacterial taxon in one site explained as much as 48% of the variation in that taxon’s growth rate in a second site (Supplementary Table 5). The amount of variation explained by each regression was the lowest when comparing the growth of ASVs between Angelo and Sedgwick, the two most distant sites in our gradient, that also have the largest difference in mean annual precipitation.

**Figure 5:**
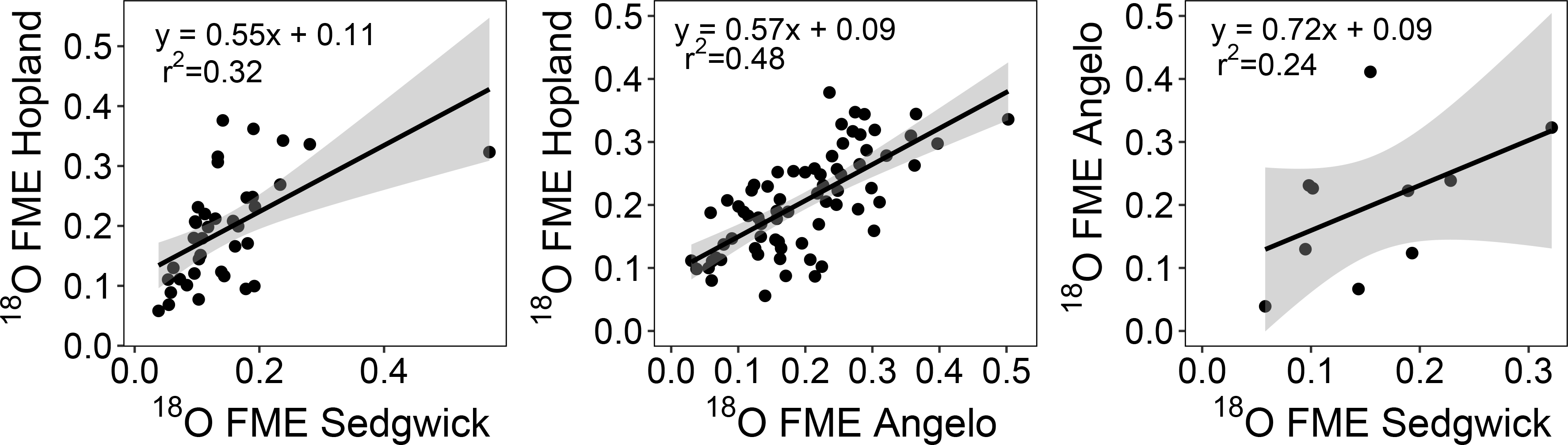
Linear regressions (black line) of fraction of maximum enrichment ^18^O (FME) of growing microbial taxa that co-occurred and grew in at least two sites from a latitudinal and precipitation gradient of annual grasslands in CA (Sedgwick>>Hopland>>Angelo). Shaded area shows 95% C.I. for the regression. Detailed regression results are presented in Supplementary Table 5. FME ^18^O is computed as the excess atom fraction ^18^O of a taxon divided by the enrichment of the soil water during the qSIP incubation.

## 4. Discussion

In this study, we used quantitative stable isotope probing (qSIP) to assess total and growing soil microbial communities across three California annual grassland ecosystems with Mediterranean climates that span a rainfall gradient. We obtained samples for this study in early 2018 during the wet season when differences in soil moisture between our sites were minimized and assessed multiple edaphic variables to characterize how variation in rainfall influences the physiochemical environment experienced by soil microorganisms at each site. We also measured the radiocarbon (^14^C) age of the soils to investigate how soil C persistence varies along the gradient and might be related to microbial growth potential. We hypothesized that actively growing communities would be more compositionally similar across the gradient than the total background microbiome due to the similarity in soil water content during the weeks preceding sampling. We also predicted that the long-term legacy effect of soil water limitation would be reflected in lower soil C, and lower growth capacity at the driest site even when soils were moisture replete.

### 4.1 Soil physiochemical environment

We characterized soils from the three California grasslands to assess how diverging rainfall regimes have shaped the physiochemical environment that microorganisms experience within each ecosystem. The three sites developed upon sedimentary parent material, primarily Franciscan sandstone and shale (Table 1). The three soils contain the same minerals but in varying proportions, pointing to the shared characteristics of parent material yet diverging weathering regimes (Supplementary Table 3). pH decreased with increasing MAP, which is likely driven by increased leaching of base cations with higher amounts of precipitation (Voroney and Heck, 2015). Some soil characteristics did not obviously correlate with rainfall patterns. Effective cation exchange capacity was highest at the driest site, Sedgwick, which is likely driven by its high clay content, but does not differ between the wet and intermediate sites. Additionally, soil carbon and nitrogen content, two important drivers of microbial composition and function, varied very little between the sites (Table 2), suggesting a similar backdrop for microbial growth, beyond the substantial differences in MAP.

### 4.2 Growing communities resemble the structure of total communities

Microbial growth drives elemental cycling in soils, so understanding the factors controlling which taxa grow is critical for soil biogeochemistry. In ecosystems with Mediterranean climates, strong seasonal changes in soil water may be an important determinant of growing community structure. These patterns have clear effects on instantaneous microbial growth rates, but may also have more pervasive, long-term legacy effects (Schimel, 2018). In this study, we characterized the total and growing soil microbial communities in three California grassland ecosystems that span a rainfall gradient during the wet season when all soils were moisture replete (Supplementary Table 2). We hypothesized that growing communities would be compositionally more similar to each other than total communities due to the convergence in environmental conditions, namely soil moisture and an actively growing plant community, during the weeks prior to sample collection. In contrast to this expectation, both growing and total communities were distinct between sites, and the distinctions between sites were of the same order of magnitude for total and growing communities (Figure 2, Supplementary Figure 3).

Microbial ecologists have long sought a reliable metric of metabolic activity with taxonomic resolution. For many years, it was assumed that rRNA extracted from environmental samples could be used to identify metabolically active microbial populations (Jones and Lennon, 2010; Campbell and Kirchman, 2013; Bowsher et al., 2019). Sequencing of rRNA has quantified strong compositional differences between active and total soil microbial communities at regional scales, and has also shown that rRNA-based soil microbial communities are more similar over space than DNA-based communities (Angel et al., 2013; Baubin et al., 2019; Locey et al., 2020). Our data suggest that a site’s climate history, rather than recent environmental conditions, place a stronger imprint on the structure of growing soil microbial populations. This is another piece of evidence suggesting that rRNA is an imperfect indicator of metabolically active populations (Blazewicz et al. 2013). While heavy-water DNA qSIP quantifies growing microbial populations, the tight coupling between DNA and RNA synthesis in soils suggests that the majority of active taxa are likely also growing (Papp et al., 2018b).

We characterized microbial communities at a time of the year when soils were replete with moisture and when environmental conditions should generally be favorable for microbial growth. However, the growing taxa in this study were still only a subset (28-58%) of all taxa detected by 16S rRNA sequencing, indicating a substantial presence of dormant organisms or relic DNA (Supplementary Table 4). While this finding is consistent with our current understanding of microbial communities in soils (the majority of microorganisms are believed to be dormant and relic DNA is estimated to comprise on average 40% of prokaryotic and fungal DNA) (Blagodatskaya and Kuzyakov, 2013; Carini et al., 2017), it raises important questions about microbial growth in these ecosystems. For example, are non-growing taxa merely the genetic remnants of dead microbes or are they perhaps dormant taxa poised for growth at another time of the year? For example, studies in California annual grasslands have documented microbial succession during the growing season of *Avena spp*., showing that different microbial taxa grow at different points throughout the wet season (Shi et al., 2015, 2016; Nuccio et al., 2020). It is also possible that the average generation time of some microbial taxa may be so slow that their growth could not be detected during the week-long incubation we conducted. These questions highlight the need for experiments clarifying the *in situ* limits and controls of growth in the soil microbiome, information that is fundamental for identifying what maintains the tremendous microbial diversity and functional potential in soils.

### 4.3 Persistent Effect of Climate on Microbial Growth

Microbial growth rate approximates a microorganism’s contributions to elemental cycling, so it is essential to understand whether the climate history of a site can influence microbial growth rates in nature. We hypothesized that a history of repeated exposure to low soil water potential at the driest site would result in lower microbial growth even when soil water was not limiting. Consistent with this expectation, community and phylum mean growth was higher at the wet and intermediate sites, Hopland and Angelo, than at the driest site, Sedgwick (Figure 3). Variation in microbial growth did not appear to be driven by pH, as we would have expected increasing substrate availability and near-neutral pH levels to, on average, facilitate faster bacterial growth rates at Sedgwick (Rousk et al., 2009). Microbial growth rates did not appear to be driven by differences in soil texture either, as clay content (which is highest at Sedgwick) is thought to support faster growth rates by providing more habitable surface area or pore space for soil microorganisms (Bååth, 1996; Uhlířová and Šantrůčková, 2003). While precipitation is the apparent causal factor for the variation in microbial growth rates we observed, we cannot entirely rule out temperature, rainfall periodicity, or plant productivity as proximal causes.

We suspect the variation in microbial growth in our soils reflects a legacy effect of the precipitation regime, particularly at our driest site, Sedgwick. Similar findings were documented for a rainfall gradient in Texas, where microbial growth decreased with lower MAP when measured under controlled laboratory conditions (Leizeaga et al., 2021). However, the literature directly testing the legacy effect of low soil moisture on growth rate is inconsistent. While a number of short-term field and lab experiments have found that historic exposure to drought reduces microbial growth rate, field experiments operating on longer time scales (>10 years) have failed to measure the same effect, possibly due to microbial acclimations to drought in the long-term studies that do not confer reduced growth rates (Meisner et al., 2013; Rousk et al., 2013; Hicks et al., 2018; Rahman et al., 2018; Nijs et al., 2019).

Historic exposure to low soil water potential can alter microbial function through shifts in community composition or by directly altering the physiology of taxa (Evans and Wallenstein, 2014). Our findings suggest that variability in microbial growth rate was driven by alterations in community composition via environmental filtering for slow growing taxa at our driest site. NRI and NTI indices show that communities at Sedgwick display phylogenetic clustering, which indicates environmental filtering at this site (Supplementary Table 6). Within the community of microbes that were growing at the driest site, most (155/197 ASVs) were found growing at this site alone, and the ASVs that were uniquely growing at Sedgwick had slower growth, on average, than taxa that occurred and grew in at least one other site (Figure 4). Thus, we infer that reduced microbial growth at our driest site appears to be influenced predominantly by relatively slow-growing taxa that are unique to this site.

Since we measured microbial growth at the ASV level using qSIP and 16S rRNA marker gene sequencing, we lack direct evidence regarding the traits and genomic potential of the microbial taxa that exhibited faster growth at our driest site. However, under the conditions evaluated here (wet season, moisture replete soils), the consistent arrangement of growth rates among taxa from site to site (Figure 5) most likely reflects the influence of genetic constraints on growth. This finding is consistent with the idea that evolutionary history influences growth rate and suggests that the phenotype of *in situ* growth rate for a given taxon spans a range constrained by its genetics and physiology, constraints that persist across rainfall gradients, edaphic properties, and biological communities (Morrissey et al., 2019). Repeated exposure to low soil water potential at Sedgwick may favor the success of microbial taxa whose physiological traits confer resistance to water stress and contribute to slow growth. While not statistically significant (P=0.33, ANOVA), the family Comamonadaceae exhibited faster in growth Sedgwick relative to the other sites (Supplementary Figure 6) and members of this family can accumulate polyhydroxyalkanoates within the cell for C storage and have genes for carotenoid biosynthesis, EPS synthesis and hydrolysis, and trehalose synthesis (De Luca et al., 2011; dos Santos et al., 2014). Future studies combining qSIP with metagenomic sequencing may help elucidate the nature of genetic constraints on microbial growth in these soils.

### 4.4 Soil C persistence (Δ^14^C) in the grassland sites

Bulk soil Δ^14^C values indicate that soil C age increased with MAP (Figure 1), consistent with findings from a recent global scale synthesis (Heckman et al., 2022). The effect of MAP on soil C persistence is driven by multiple factors that can vary by soil fraction and depth (Heckman et al., 2022). Recent models of soil C cycling emphasize physiochemical stabilization mechanisms and microbial necromass as a precursor for persistent soil C (Lavallee et al., 2020). In our study, microbial growth, and presumably necromass production, also increased with increased MAP (Figure 3). If necromass-C is preferentially stabilized in these soils, this could result in more persistent soil C over time, leading to an older average age for bulk soil C. In addition, greater rates of vertical flushing of water at our highest MAP site could also facilitate weathering and the development of secondary clay, aluminum, and iron oxide minerals that are key contributors to soil C storage and persistence (Rasmussen et al., 2018; Slessarev et al., 2022).

Our supposition that soil C is older in the wetter sites due to the preferential stabilization of necromass-C is consistent with current literature on soil C cycling in grasslands. The transformation of plant-derived substrates to microbial products is believed to be a key process in the stabilization of soil C (Cotrufo et al., 2012; Kallenbach et al., 2016) and appears to have a pronounced importance in grasslands (Cotrufo et al., 2012; Kallenbach et al., 2016; Angst et al., 2021). In grassland ecosystems, soil C age is linked to SOC chemistry such that products that are more microbial in nature tend to be older (Heckman et al., 2021). Soil C:N, the only indicator of soil C quality in this study, did not vary between our sites (Table 2). However, soil C:N is a broad proxy and a more targeted molecular characterization of soil C is needed to conclude whether microbial processing of C increases with MAP along the gradient studied here.

## 5. Conclusions

Soil water is an important controller of microbial growth and Earth system models project significant changes in the timing and intensity of precipitation events globally (Dore, 2005; Schimel, 2018). Seasonal changes in soil moisture have clear effects on instantaneous microbial growth rates and our study demonstrates that soil water also has a more pervasive, long-term legacy effect on microorganisms. Our data show that a site’s historical climate regime, rather than recent environmental conditions, strongly influences the structure of growing microbial communities. Our study also demonstrates that a history of exposure to low soil moisture reduces microbial growth, likely through shifts in community composition. Since changes in microbial community structure operate over long time scales, ecosystems may experience a pervasive legacy effect of climate history as precipitation regimes change. Lastly, our study demonstrates that taxon-specific growth rates are remarkably consistent between three grassland sites from a wide precipitation gradient, consistent with the idea that evolutionary history influences microbial growth and points to the genetic and physiological constraints on growth rate in nature.

## Acknowledgements

We thank Erin Nuccio, Amrita Bhattacharyya, Christina Ramon, and Aaron Chew for assistance with field sampling and lab analyses; John Bailey and Allison Smith at Hopland Research and Extension Center for help gathering meteorological data; and the Oregon State University Soil Lab for chemical and physical soil analyses. We would like to thank Thomas Bristow, at the NASA Ames Research Center, for allowing access to his lab space for qXRD analysis, and the Stable Isotope Laboratory at UC Berkeley for stable isotope analyses. MM Foley was supported by a National Science Foundation Graduate Research Fellowship while completing this work.

This research was supported by the U.S. Department of Energy, Office of Biological and Environmental Research, Genomic Science Program ‘Microbes Persist’ Scientific Focus Area (#SCW1632) at Lawrence Livermore National Laboratory (LLNL) and a subcontract to Northern Arizona University. Work conducted at LLNL was conducted under the auspices of the US Department of Energy under Contract DE-AC52-07NA27344.

## Notes

### Competing Interest Statement

The authors have declared no competing interest.

