## Supplementary Information for "Active populations and growth of soil microorganisms are framed by mean annual precipitation in three California annual grasslands"

**Supplementary Table 1** Air temperature and precipitation data for three northern CA annual grassland sites. Table includes weather data for the calendar year and the 30-day period leading up to sample collection. Data for Angelo and Sedgwick were gathered from the Western Regional Climate Center (https://wrcc.dri.edu/). Data for Hopland were gathered via personal communication with Allison Smith at Hopland Research and Extension Center.

|  | Avg. Air Temp (max/min°C)  Year Prior | Cumulative precipitation (mm)  Year Prior | Avg. Air Temp (°C)  30-days prior | Cumulative precipitation (mm)  30-days prior |
| --- | --- | --- | --- | --- |
| Angelo | 26/0 | 2446 | 3.76 | 206 |
| Hopland | 28/-3 | 660 | 8.76 | 155 |
| Sedgwick | 32/5 | 434 | 10.35 | 137 |

**Supplementary Table 2** Gravimetric soil moisture upon collection from the field and during the qSIP incubation (mean ± standard error, n = 3). Enrichment of the soil water during qSIP incubation (mean ± standard error, n = 3) was estimated based on the mass of 98.15 at% ^18^O-H_2_O added to soils and mass of natural abundance water present in sample at the start of the incubation.

|  | Field moisture (%)  Avg ± SE | qSIP Incubation moisture (%)  Avg ± SE | Soil water enrichment (at %)  Avg ± SE |
| --- | --- | --- | --- |
| Angelo | 23 ± 1 | 28 ± 1 | 87 ± 4 |
| Hopland | 17 ± 1 | 24 ± 0.3 | 94 ± 1 |
| Sedgwick | 25 ± 1 | 28 ± 1 | 87 ± 2 |

**Supplementary Table** **3** Relative abundance of soil minerals measured by quantitative X-ray diffraction (mean ± standard error, n = 3).

|  |  | Angelo (wt %)  Avg ± SE | Hopland (wt %)  Avg ± SE | Sedgwick (wt %)  Avg ± SE | |
| --- | --- | --- | --- | --- | --- |
| Non-Clay | Quartz | 32.33 ± 2.46 | 40.40 ± 0.97 | | 32.76 ± 0.87 |
|  | Plagioclase | 21.41 ± 2.81 | 18.63 ± 0.87 | | 19.04 ± 0.54 |
|  | K-feldspar | 4.76 ± 0.80 | 2.82 ± 0.22 | | 5.58 ± 0.82 |
| Clay | Muscovite | 22.81 ± 0.64 | 19.27 ± 2.59 | | 11.74 ± 0.94 |
|  | Chlorite | 13.03 ± 2.60 | 10.38 ± 0.41 | | 14.67 ± 3.35 |
|  | Kaolinite | 5.66 ± 2.85 | 8.50 ± 2.74 | | 16.21 ± 3.04 |

**Supplementary Table 4** The percent of the total microbial community at each site that was quantified as growing. The total number of ASVs represents the number of unique ASVs summed across replicates (n=3) at each site inferred from 16S rRNA sequencing. The number of ASVs growing is the number of taxa with 95% confidence intervals for excess atom fraction ^18^O (EAF) significantly greater than zero at each site. The proportion of the community growing was calculated as the number of ASVs growing divided by the total number of ASVs at a site.

|  | Total No. ASVs | No. ASVs growing | Proportion of community growing |
| --- | --- | --- | --- |
| Angelo | 510 | 141 | 28% |
| Hopland | 590 | 282 | 48% |
| Sedgwick | 339 | 197 | 58% |

**Supplementary Table 5** Summary statistics of linear regressions of an ASV’s fraction of maximum enrichment ^18^O (FME) in one site versus the FME ^18^O of that same ASV in another site, as shown in Figure 5. FME ^18^O is computed as the excess atom fraction ^18^O of a taxon divided by the enrichment of the soil water during the qSIP incubation. Bold values indicate p<0.05.

| Model | Slope estimate | Slope standard error | P-value | Residual Std Error | R^2^ |
| --- | --- | --- | --- | --- | --- |
| Hopland ~ Sedgwick | 0.55 | 0.14 | **<0.01** | 0.07 | 0.32 |
| Hopland ~ Angelo | 0.57 | 0.07 | **<0.01** | 0.06 | 0.48 |
| Sedgwick ~ Angelo | 0.72 | 0.45 | 0.15 | 0.11 | 0.24 |

**Supplementary Table 6** Results from NRI and NTI analyses. *N* is the number of taxa in community, *NRI* is net relatedness index, and *NTI is* nearest taxa index. Each metric was calculated for individual soil cores (n=3). Bold values indicate p<0.05. Significantly positive NRI and NTI values above 1.96 indicate phylogenetic clustering.

|  | Replicate | N | NRI | NTI |
| --- | --- | --- | --- | --- |
| Angelo | 1 | 189 | **2.76** | **2.38** |
|  | 2 | 337 | **2.55** | **1.93** |
|  | 3 | 297 | **2.76** | -0.27 |
| Hopland | 1 | 337 | 1.56 | **1.94** |
|  | 2 | 267 | **2.91** | 0.29 |
|  | 3 | 324 | **1.94** | -0.83 |
| Sedgwick | 1 | 204 | **2.19** | **2.88** |
|  | 2 | 192 | **3.14** | **3.58** |
|  | 3 | 211 | **2.38** | 1.53 |

**Supplementary Table 7** Results from phylogenetic signal analysis of taxon-specific excess atom fraction ^18^O (EAF). For both Pagel’s λ and Blomberg’s K, values approaching 1 indicate a phylogenetic signal. Bold values indicate P < 0.05.

|  | Pagel’s λ | Blomberg’s K |
| --- | --- | --- |
| Angelo | **0.82** | **0.30** |
| Hopland | **0.85** | **0.10** |
| Sedgwick | **0.68** | **0.10** |

**Supplementary Figure 1** Precipitation and temperature data for our three northern CA annual grassland sites from 2017-2019. Grey bars show monthly cumulative precipitation while red points show monthly average air temperature. Black asterisks denote the month when samples were collected. Data for Angelo and Sedgwick were gathered from the Western Regional Climate Center (https://wrcc.dri.edu/). Data for Hopland were gathered via personal communication with Allison Smith at Hopland Research and Extension Center.

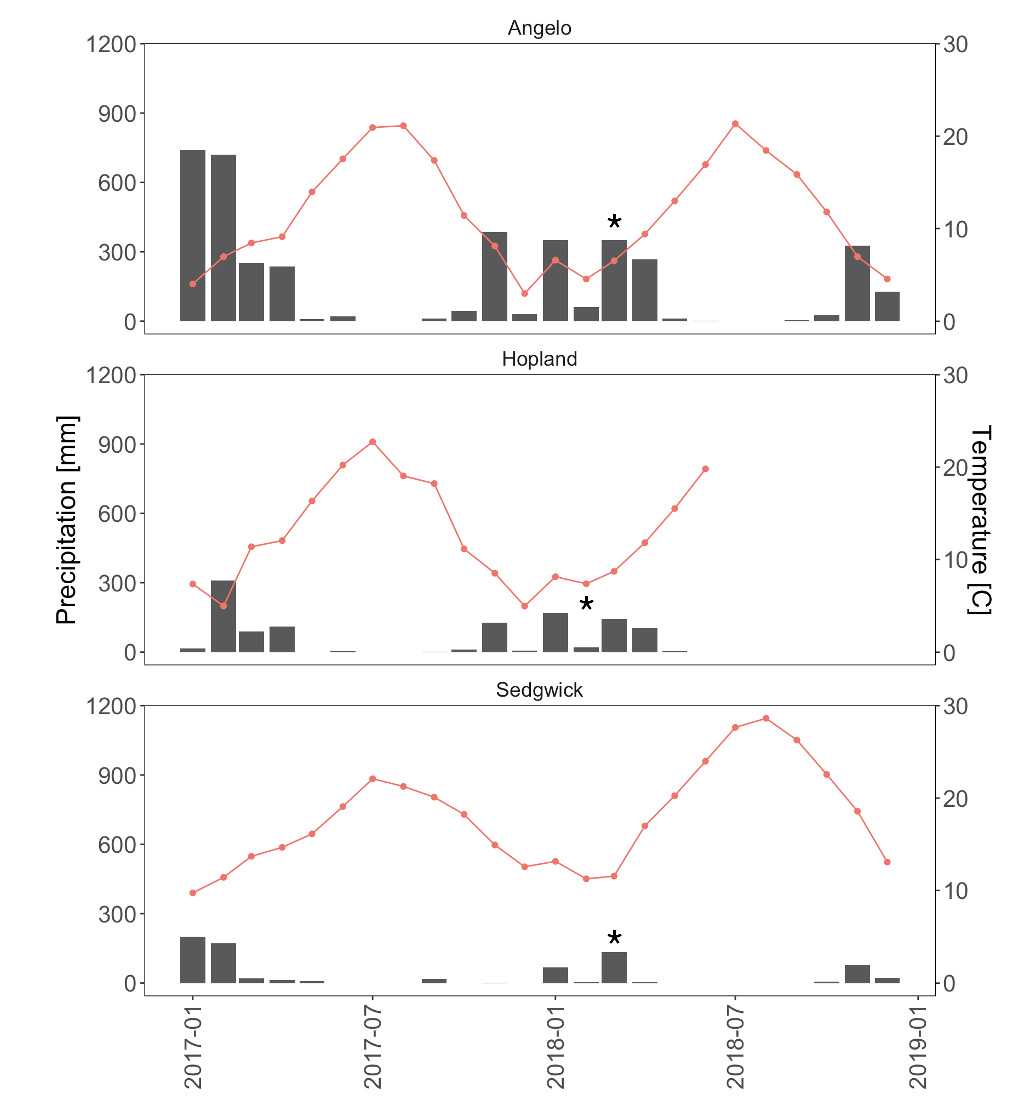

**Supplementary Figure 2** Soil water retention curves for our three northern CA annual grassland sites.

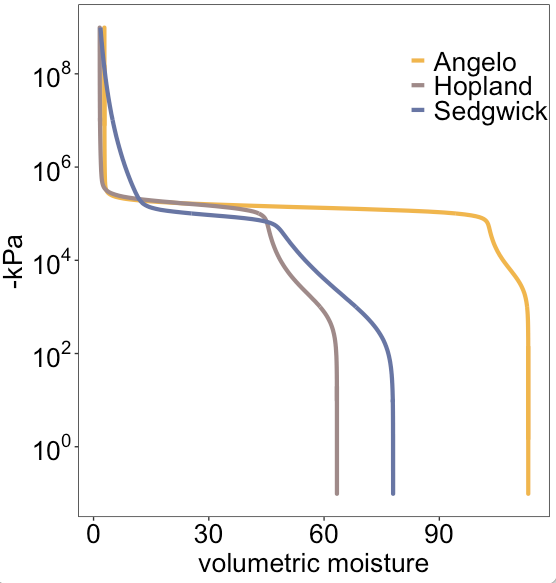

**Supplementary Figure 3** Mean pairwise dissimilarity between sites for the present microbial community and growing microbial community. Dissimilarity did not differ for growing versus total communities when comparing between Angelo and Hopland (t-test, p = 0.95), Angelo and Sedgwick (t-test, p = 0.1699), nor Hopland and Sedgwick (t-test, p = 0.6453). Bars show 95% confidence intervals for the means.

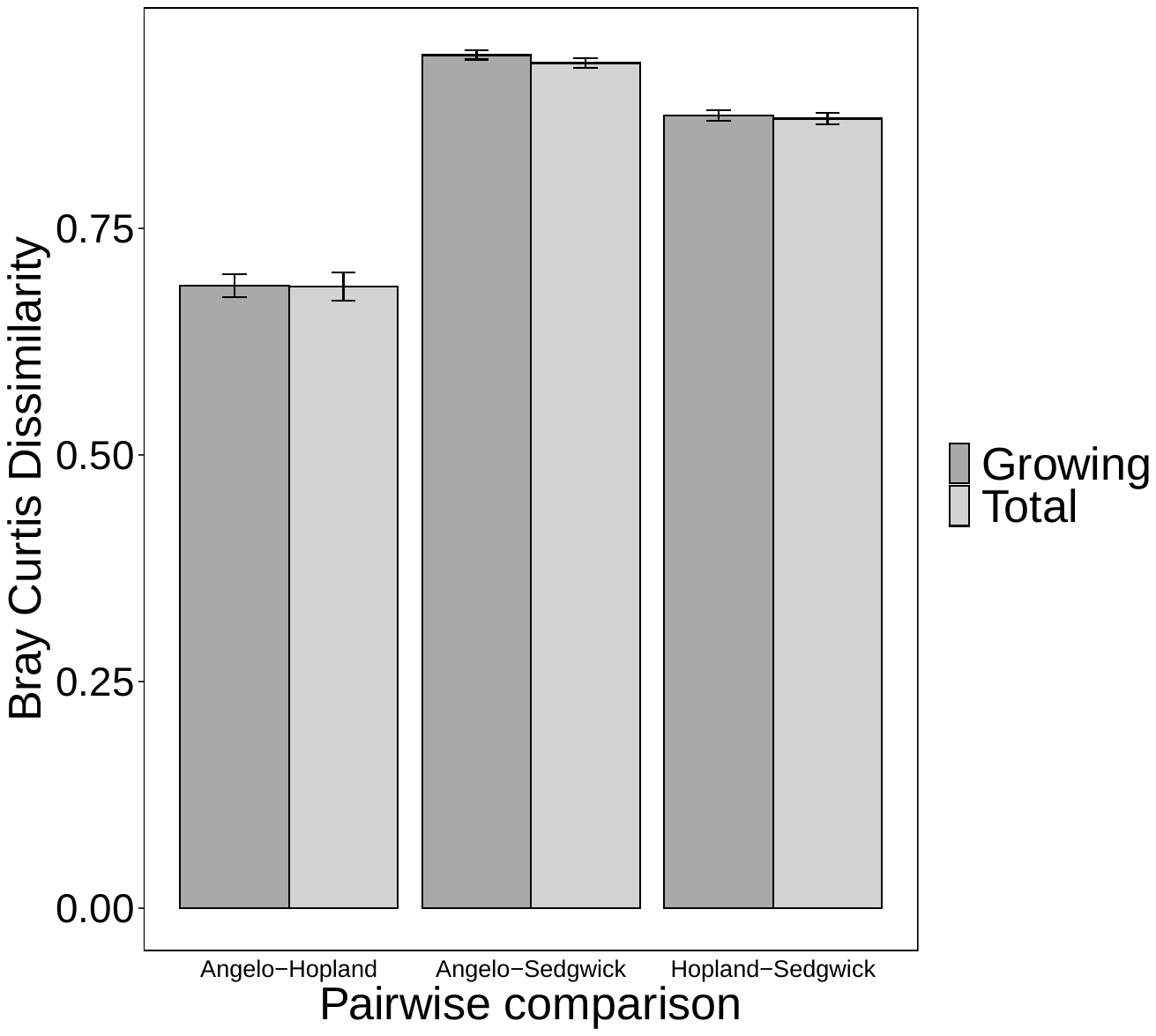

**Supplementary Figure 4** Relative abundances of phyla in A) total and B) growing communities at Angelo, Hopland, and Sedgwick.

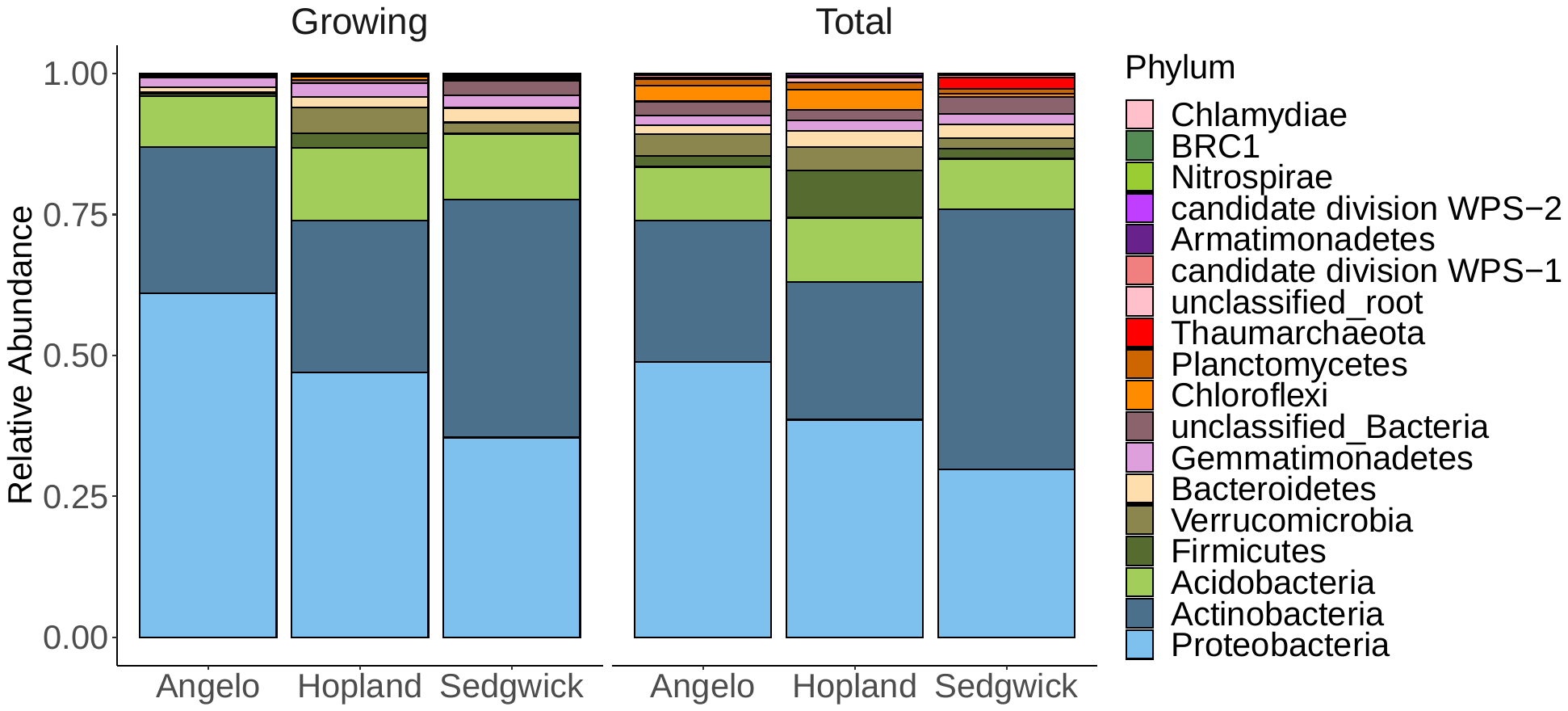

**Supplementary Figure 5** Fraction of maximum enrichment ^18^O (FME) of a taxon versus its relative abundance. FME ^18^O is computed as the excess atom fraction ^18^O of a taxon divided by the enrichment of the soil water during the qSIP incubation. Each point represents an ASV whose enrichment was measured via qSIP and its relative abundance was assessed through sequencing of unfractionated DNA extracted from soils at the grassland sites. Values for relative abundance are log-transformed.

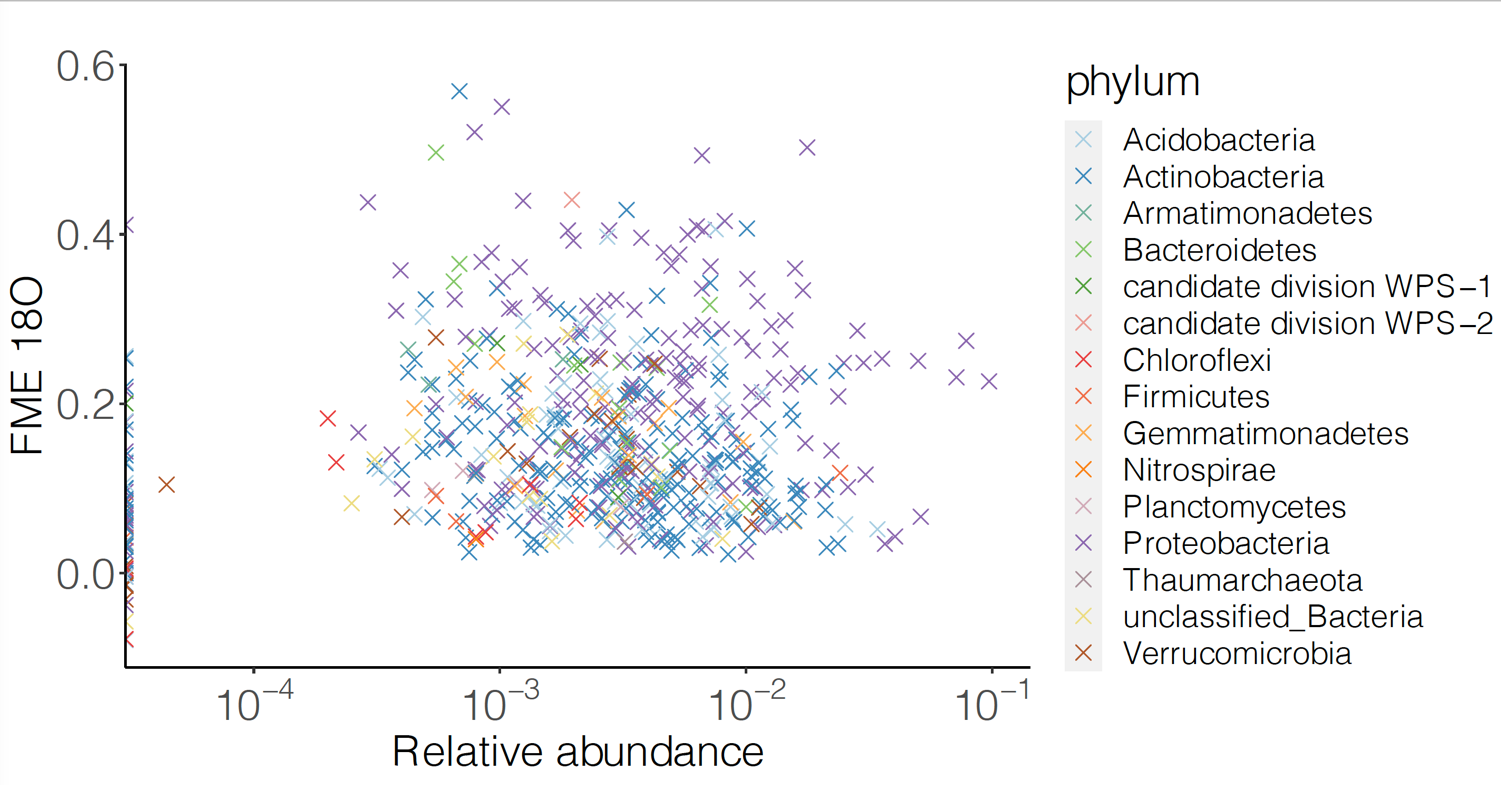

**Supplementary Figure 6** Fraction maximum enrichment ^18^O (FME) of microbial families. FME ^18^O is computed as the excess atom fraction ^18^O of a taxon divided by the enrichment of the soil water during the qSIP incubation. Points represent mean ^18^O FME of a family and bars indicate standard deviation. Letters indicate significant differences between sites (p<0.05, Fisher’s LSD).

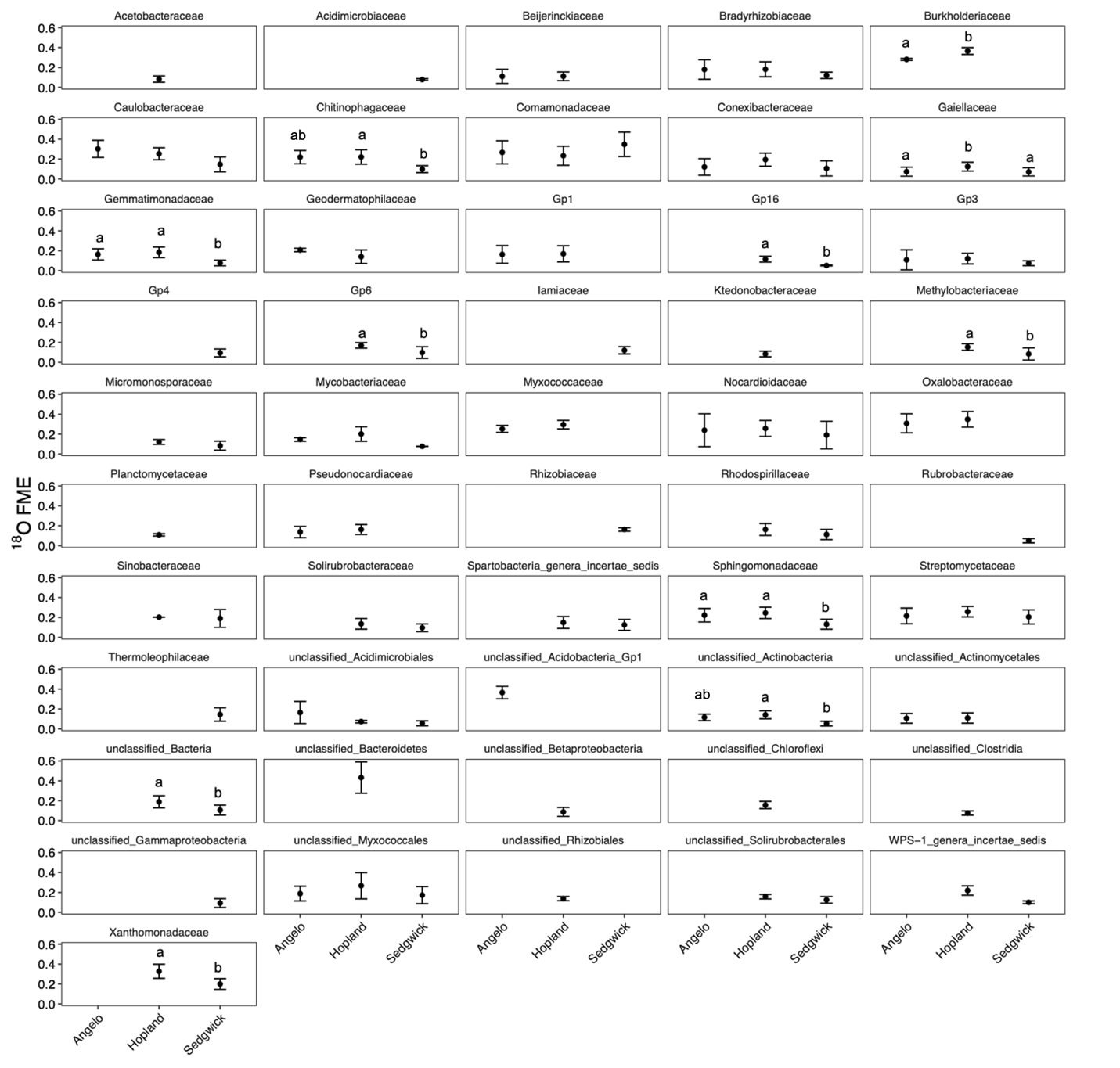
